# Differences in homologous recombination and maintenance of heteropolyploidy between *Haloferax volcanii* and *Haloferax mediterranei*

**DOI:** 10.1101/2022.03.26.485934

**Authors:** Ambika Dattani, Itai Sharon, Ella Shtifman-Segal, Shachar Robinzon, Uri Gophna, Thorsten Allers, Neta Altman

## Abstract

Polyploidy, the phenomenon of having more than one copy of the genome in an organism, is common among haloarchaea. While providing short-term benefits for DNA repair, polyploidy is generally regarded as an ‘evolutionary trap’ that by the notion of the Muller’s ratchet will inevitably conclude in the species’ decline or even extinction due to a gradual reduction in fitness. In most reported cases of polyploidy in archaea, the genetic state of the organism is considered as homoploidy i.e. all copies of the genome are identical. Here we demonstrate that while this is indeed the prevalent genetic status in the halophilic archaeon *H*. volcanii, its close relative *H. mediterranei* maintains a prolonged heteroploidy state in a non-selective environment once a second allele is introduced. Moreover, a strong genetic linkage was observed between two distant loci in *H. mediterranei* indicating a low rate of homologous recombination while almost no such linkage was shown in *H. volcanii* indicating a high rate of recombination in the latter species.

We suggest that *H. volcanii* escapes Muller’s ratchet by means of an effective chromosome-equalizing gene-conversion mechanism facilitated by highly active homologous recombination, whereas *H. mediterranei* must elude the ratchet via a different, yet to be elucidated mechanism.

## Introduction

Polyploidy, having multiple copies of the same chromosome in a cell, is a well-known property of different haloarchaeal species and was demonstrated as well as characterized in *H. volcanii. H. mediterranei* and *H. salinarum*. (Breuert *et al*. 2006). Polyploidy is also found in multiple additional members of the Euryarcheota (Ludt and Soppa 2019). The number of chromosomal copies a cell harbors depends on the growth phase of the culture and vary from 1 (Malandrin *et al*. 1999) and up to 55 (Spaans *et al*. 2015). Potential evolutionary benefits of polyploidy vary from one species to another and include use of DNA as a phosphate storage (Zerulla *et al*. 2014), and an improved ability to repair DNA breaks (Hildenbrand *et al*. 2011) (Perez-Arnaiz *et al*. 2020), including those generated by autoimmunity caused by the microbial immune system CRISPR-Cas (Stachler *et al*. 2017).

Homologous recombination is facilitated by polyploidy since it guarantees multiple wild type templates that could be used for double strain break repair in archaea (Jones and Baxter 2017) (Kish and DiRuggiero 2008) and bacteria (Zahradka *et al*. 2006). Polyploidy however could also be costly, especially in harsh mutagenic environments. Gene conversion and equalization of the genome copies (Soppa 2011) as well as lateral gene transfer and homologous recombination (Papke *et al*. 2004) was suggested as possible strategies to overcome the accumulation of deleterious alleles and a way to lessen the mutation load (Kondrashov 1994) and escape genetic degradation via Muller’s ratchet (Markov and Kaznacheev 2016) in those asexual non-mitotic organisms.

While polyploidy is common in many prokaryotes (archaea and some bacteria) (Markov and Kaznacheev 2016) (Soppa 2011), those polyploids are generally regarded as homopolyploid, namely having genome copies that contain the same genetic information. This contrasts with the prevalent polyploidy in plants that display autopolyploidy (a multiplication of the chromosomal set) in some species alongside allopolyploidy (chromosome doubling accompanied by hybridization of two parents, often of different species) in others (Stuessy and Weiss-Schneeweiss 2019) (Soltis *et al*. 2015).

In the absence of mitosis, polyploidy could lead to an accumulation of deleterious mutations, and an ongoing reduction in the fitness of the population, unless there are mechanisms that can counter this process (Markov and Kaznacheev 2016) (Soppa 2011). One such mechanism could be lateral gene (or in this case allele) transfer between cells and indeed *H. volcanii* and *H. mediterranei* are known in their ability to create long lasting cytoplasmic bridges that enable the passage of DNA within the species (Rosenshine *et al*. 1989) and even creating a hetero-diploid fused interspecies cell hybrid (Naor *et al*. 2012). These haloarchaea thus somewhat resemble plant allopolyploid species. These heteropolyploid hybrids are possible to maintain under selective pressure but are quickly resolved back to homopolyploidy in the absence of such selection. While it is well established that the application of selective pressure on *H. volcanii* can preserve a heteropolyploid state created by cell-cell mating or an artificial semi-heteropolyploidy created via plasmid transformation (Lange *et al*. 2011), those so-called merodiploid states are lost when the selective pressure is relieved and the genome returns to its homoploid state.

When considering the polyploidy of *H. volcanii*, the ease in which genomic deletions and mutations are introduced into this species is quite surprising (Bitan-Banin *et al*. 2003) (Allers and Mevarech 2005). A genome-homogenizing process that agrees with the observations discussed above, seems to underlie the relative ease of creating gene knock-outs in *H. volcanii* despite its large genome copy number. Similar observations have been made in the methanogen *Methanococcus maripaludis (Hildenbrand et al. 2011*). In contrast, we have observed that when using the same vectors and procedures commonly applied for making gene knockouts for *H. volcanii, H. mediterranei* gene knockouts require laborious passages in selective media, even under low phosphate conditions reduced where ploidy is (Turgeman-Grott personal communication). This observation suggests that while *H. volcanii* utilizes a genome homogenizing mechanism, *H. mediterranei* does not homogenize its genome copies as efficiently as *H. volcanii* does. This suggests that the two related species might also demonstrate additional differences in replication and DNA repair.

The ability of a polyploid cell to equalize quickly up to 20 copies of its genome under selective pressure could be explained either by segregation and a strong heteropolyploidy fitness cost, or by a robust homologous recombination and gene conversion mechanism (Hildenbrand *et al*. 2011); the latter are presumably also important for additional cellular processes (White and Allers 2018) (Barilla 2016). In this paper we have studied the ability of both *H. volcanii* and *H. mediterranei* to retain a heteropolyploid state artificially created by cell-cell fusion, when selection is relieved. We also examine their respective subsequent recombination rates in the absence of selection. We show that when selection is relieved, *H. volcanii* equalizes its multiple genome copies efficiently while *H. mediterranei* does not. We also show surprising differences in recombination rates between the two species, using both cell-cell mating and plasmid-chromosome recombination assays. While spontaneous homologous recombination in *H. volcanii* is extremely common, a strong genetic linkage is observed between two distant loci in *H. mediterranei*. Comparative RNA-seq displayed no obvious differences in expression levels of factors of the DNA replication pathway. A significant difference in expression levels of factors of the homologous recombination pathway was observed in stationary-phase *H. mediterranei* cells, with an increase in the recombination mediator *radB* expression.

## Results

### The heteropolyploid state is stable under selection

To investigate differences in genome-homogenization and recombination between *H. mediterranei* and *H. volcanii*. we generated two pairs of strains, an *H. volcanii* pair and an *H. mediterranei* pair, to be employed in mating assays (either *volcanii-volcanii* or *mediterranei-mediterranei* mating). The *H. volcanii* paternal strain (Allers *et al*. 2004) contained deletions in the *trpA* and *hdrB* genes (tryptophan and thymidine auxotrophy respectively) and the two strains were constructed such that the *trpA* or *hdrB* cassettes were inserted back into the same ectopic location, instead of the ASC-like gene HVO1585, which was previously demonstrated to be a non-essential ORF (Moshe Mevarech personal communication). This genetic background allowed for the selection of mated cells on growth medium lacking tryptophan and thymidine. Typically, such mating assays result in a homopolyploid cell containing a genome harboring both cassettes, which would be the outcome of recombination events. The unique construction of the current mated pair however prevents such recombination events from happening as the cassettes were inserted into the same genomic location, hence the mated cells are forced to contain two different types of genomes and are essentially held under selection in a heteropolyploid state. In addition to the metabolic selectable marker, a second marker was introduced to one of the pair strains – a deletion in the *crtI* (HVO_2528) gene, which is involved in biosynthesis of the carotenoid pigment bacterioruberin and is not essential nor has an effect on fitness (Turkowyd *et al*. 2020). This deletion resulted in a white phenotype (WR807) in contrast with the other strain that remained naturally pigmented (red), (WR780). Essentially the same process was repeated with *H. mediterranei* but for practical reasons the *pyrE* gene (HFX_0318, uracil auxotrophy) was used as a selective marker instead of *folA (hdrB* homologue) to generate strain UG482 (white), along with *trpA* (HFX_0748) to generate strain UG480 (pink-red, see Methods). In total, two pairs of strains were constructed, each of which contained two different markers: one for auxotrophy (*H. volcanii* tryptophan/thymidine, *H. mediterranei* tryptophan/uracil), and another for color (red/white) as described in Table 1. *H. volcanii* and *H. mediterranei* heteropolyploid cells were generated by mating WR780 with WR807 and UG480 with UG482 respectively. The heteropolyploid state of the mating products was confirmed by PCR amplification of the loci that were under selection, using the primers T36+T37 and T28+T29 (Table 4).

**Table 1.** A) A summary of the strains used in this research, their genotypes, and phenotypes. *crtI* of *H. volcanii* and *H. mediterranei* are homologous. HVO_1585 is homologous to HFX_1643 B) chromosomal locations of all genetic markers used

| Strain | Specie | Genotype (partially described) | Relevant properties | Source |
| --- | --- | --- | --- | --- |
| WR780 | *H. volcanii* | $\Delta trpA$, $\Delta hdrB$, 1585::*trpA* | Thymidine auxotrophy, red | This study |
| WR807 | *H. volcanii* | $\Delta trpA$, $\Delta hdrB$, $\Delta crtI$, 1585::*hdrB* | Tryptophan auxotrophy, white | This study |
| H26 | *H. volcanii* | $\Delta pyrE2$ | Uracil auxotrophy | (ALLERS *et al.* 2004) |
| UG480 | *H. mediterranei* | $\Delta pyrE$, $\Delta trpA$, 1643::*trpA* | Uracil auxotrophy, red | This study |
| UG482 | *H. mediterranei* | $\Delta pyrE$, $\Delta trpA$, $\Delta crtI$, 1643::*pyrE* | Tryptophane auxotrophy, white | This study |
| WR505 | *H. mediterranei* | $\Delta pyrE2$ | Uracil auxotrophy | (NAOR *et al.* 2013) |

| *Haloferax volcanii* |  |  | *Haloferax mediterranei* |  |  |
| --- | --- | --- | --- | --- | --- |
| Gene name | HVO | Location on chromosome | Gene name | HFX | Location on chromosome |
| *trpA* | HVO_0789 | 708729-709562 | *trpA* | HFX_0748 | 704395-705228 |
| *pyrE2* | HVO_0333 | 301751-302281 | *pyrE* | HFX_0318 | 299911-300411 |
| *hdrB* | HVO_2919 | 2754021-2754605 | Not used |  |  |
| *ASC-like* | HVO_1585 | 1455006-1457012 | *ACS_like* | HFX_1643 | 1566531-1568528 |
| *crtI* | HVO_2528 | 2394228-2395730 | *crtI* | HFX_2250 | 2093437-2094939 |

### *H. volcanii* and *H. mediterranei* lose heterozygosity at different rates under non-selective conditions

The heteropolyploid state was stable as long as the mating products were kept under selection but started decreasing at different rates when selection was relieved as described next. To explore heterozygosity maintenance in the absence of selective pressure, single heteropolyploid *H. volcanii* or *H. mediterranei* colonies were picked into non-selective YPC liquid media and samples were taken 48h (5 biological replicates) and one week (5 additional biological replicates) after inoculation (fresh growth medium was added every other day) and were plated on non-selective media. 50 colonies were then taken from each plate and streaked on two selective media types each with selection to only one of the metabolic markers. The number of colonies that grew on both types of media was enumerated and used to calculate the percentage of heterozygous colonies in the population. To corroborate the results of these viable counts, the genotype of about 20% of the colonies was also determined by PCR and was found to match the growth phenotype (Fig 1S). We observed near-complete loss of heterozygosity in *H. volcanii*, but not in *H. mediterranei* after 1 week of growth.

After 48h in non-selective media, both *H. volcanii* and *H. mediterranei* mating products showed a decrease of the heteropolyploid state to an average of approximately 30% of screened colonies. The replicates showed a wide variation of the heteropolyploid state ranging from 4% to 56% in *H. mediterranei* and from 8% to 48% in *H. volcanii*. However, a near-complete loss of heterozygosity in *H. volcanii*, but not *H. mediterranei* was observed after one week, at which point *H. mediterranei* was still showing an approximately 30% heteropolyploidy (ranging from 6% to 88%) while 4 of the 6 *H. volcanii* replicates showed no heteropolyploidy and the other 2 replicates showed 4% and 10% heteropolyploidy (Table 2).

**Table 2.**

| **<u>Species</u>** | **<u>% heteroploids</u>** |  |
| --- | --- | --- |
|  | <u>48 Hours</u> | <u>1 week</u> |
| ***H. volcanii*** | 28±13 | 2±2 |
| ***H. mediterranei*** | 28±17 | 34±26 |

### Recombination rate is much higher in *H. volcanii* than in *H. mediterranei*

Since each chromosome of the heteropolyploid cells described above contained one color marker and one selective marker, there are four possible genotype combinations that can, in theory, exist in selected cells. Notably, since only two of these four combinations are present in any of the original strains of each species, emergence of the two other combinations can only result from recombination events in cells that contain both types of the original chromosomes.

Immediately after mating, mated cells are heteropolyploid, yet once selection is relaxed cells can segregate to a homopolyploid state (as above). We next examined the recombination rate of segregated cells in the two species. Since each strain has a unique combination of markers, it was possible to determine whether segregated (homopolyploid) cells were similar to one of the original strains or contained a new combination of color and selective marker. The colonies obtained after 48h and inferred to be homopolyploid in the previous analysis were thus analyzed (Table 3). *H. volcanii* colonies demonstrated a high rate of recombination where about 50% of the colonies retained the original (parental) marker combination while the other 50% showed a recombinant genome. In contrast, about 80% of *H. mediterranei* cells retained the original (parental) markers combination. These results suggest that in *H. volcanii* the recombination rate is higher than in *H*. *mediterranei*, especially taking into account that the genomic distance between the markers used is double in *H. volcanii* compared to *H. mediterranei* (^~^0.5M bp in *H. mediterranei* and ^~^1M bp in *H. volcanii*). Colony color-based results were validated by PCR (conducted on a sample of 5% of the colonies) and confirmed in 100% of cases.

**Table 3.**

| **<u>Species</u>** | **<u>% similarity to parents</u>** |  |
| --- | --- | --- |
|  | <u>48 Hours</u> | <u>1 week</u> |
| ***H. volcanii*** | 52±15 | 50±1 |
| ***H. mediterranei*** | 80±15 | 78±26 |

To directly compare the recombination rates of these species, we then performed plasmid-chromosome recombination (integration or “pop-in”) experiments in which *H. mediterranei* and *H. volcanii* cells were transformed with a non-replicating plasmid bearing a *pyrE2* selectable marker. In such experiments, only cells that integrate the plasmid into their chromosome via homologous recombination can grow. Four plasmids for *H. mediterranei* (pTA2684, pTA2688, pTA2692, pTA2696) and four plasmids for *H. volcanii* (pTA2686, pTA2690, pTA2694, pTA2698) were constructed for this purpose. Each such plasmid carries a DNA fragment of approximately 1000 bp that corresponds to a portion of either the *H. mediterranei* or *H. volcanii* genome. Orthologous genes were chosen for *H. mediterranei* and *H. volcanii* since they occupy similar positions in their respective genomes and share high gene similarity between species. All plasmids carry the *pyrE2* selectable marker (*pyrE2* from *H. volcanii*) that is suitable for both organisms (Table 5). Recombination rates were calculated as the percentage of the cells that integrated the plasmids compared to the viable count. Transformation efficiency was calculated using a replicating plasmid (pTA245), to control for the effect of transformation efficiency. We observed that for all DNA fragments, the plasmid integration rates were generally over 5-fold higher in *H. volcanii* than in *H. mediterranei*. (Figure 1). Taken together with the segregation experiments described above, these findings indicate significantly higher recombination rates in *H. volcanii* compared to *H. mediterranei*.

**Figure 1.**
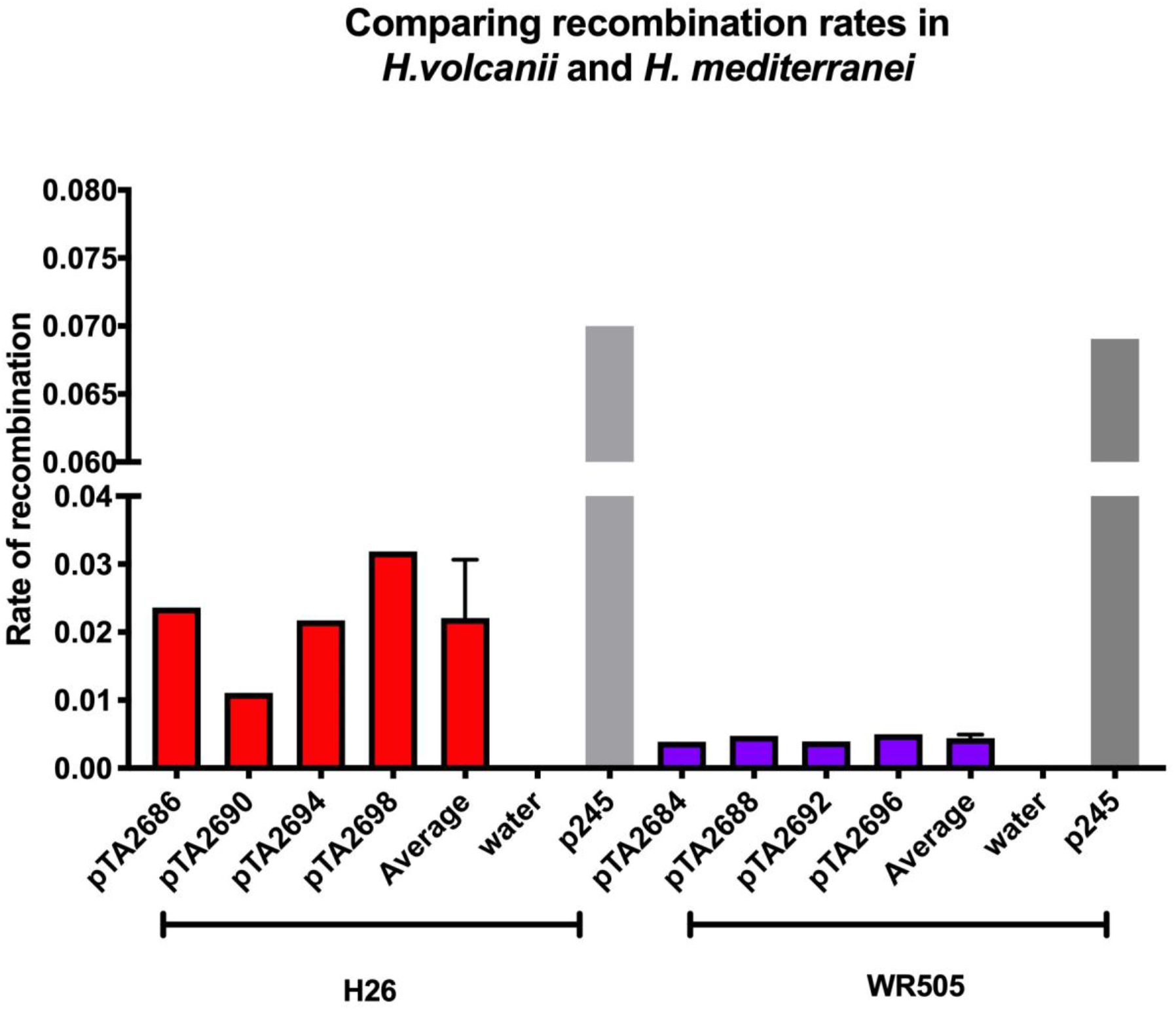

### Comparative RNA-seq analysis of the DNA replication and homologous recombination pathways in H. mediterranei and H. volcanii

To determine whether a difference in RNA expression levels could explain the variations described above, a comparative analysis of DNA replication and homologous recombination factors in both *Haloferax* species was carried out (Supplementary Table 1). Most genes involved in DNA replication and replication-restart showed no statistically-significant differences in expression levels between the two species. In terms of the homologous recombination pathway, although *radA, mre11, rad50*, and genes encoding DNA ligases had similar expression levels in both species, an increase in *radB* (Wardell *et al*. 2017) expression was noted in *H. mediterranei* stationary phase cells. This difference could result in divergent processing of DNA breaks and explain the decreased homologous recombination rate in *H. mediterranei*.

## Discussion

*H. volcanii* and *H. mediterranei* are two closely related halophilic archaea species. They share an average nucleotide sequence identity of 86.6%, are able to perform intra-species cell-cell fusion followed by an exchange of genomic fragments via recombination, and have a similar genome architecture (Naor *et al*. 2012). Despite their similarities, the two species demonstrate a variety of genetic differences, one of which is discussed in this work. We show that while *H. volcanii* seems to equalize its genome copies quickly and effectively, *H. mediterranei* can maintain a state of semi heterozygosity/heteropolyploidy for many generations; this corresponds to a high frequency of HR events in *H. volcanii*, and much lower rates in *H. mediterranei*.

Two possible mechanisms could underlie these differences in heterozygosity; modifications in chromosome segregation or variations in gene conversion and recombination mechanisms. The knowledge of archaeal chromosome segregation is limited and sparse. While it was suggested that the same SegAB dependent mechanism may apply in monoploid or diploid archaea of both Euryarchaeota and Crenarchaeota (Barilla 2016), little is known about the mechanism of chromosome segregation in polyploid archaea. Importantly, most Euryarchaeota and halophilic archaea in particular are highly polyploid (Chant and Dennis 1986) (Liu *et al*. 2013) (*Breuert et al*. 2006) (Spaans *et al*. 2015) (Barilla 2016) (Zerulla and Soppa 2014) (Ludt and Soppa 2019), further complicating segregation issues.

While not much is known about differences in chromosome segregation between the two species, evidence about differences in the way that recombination is connected to replication does exist. An *H. volcanii* strain lacking all four origins of replication was strictly dependent on the RadA recombinase, hence it was proposed that recombination alone can support DNA replication in *H volcanii*. This phenomenon was also later suggested for the hyperthermophilic archaeon *T. kodakarensis*, which is also polyploid (Gehring *et al*. 2017). In contrast, an *H. mediterranei* strain that had its three major chromosomal origins deleted was viable but slow growing and was shown to be dependent on a dormant origin (*oriC4*) that became essential in the absence of the other three. Furthermore, unlike the origin-deleted mutant of *H. volcanii*, that *H. mediterranei* mutant showed higher *radA* expression but was viable in a *radA* suppressed background, indicating that it was dependent on the cryptic origin rather than on HR for replication. These past studies indicate that in the total absence of origins, *H. volcanii* can depend on recombination-initiated replication whereas *H. mediterranei* cannot. This could be attributed either to differences in genome replication or a failure of *H. mediterranei* HR machinery to support DNA replication.

Our observations that the recombination frequency appears to be much lower in *H. mediterranei* than in *H. volcanii* suggests that homologous recombination operates at a different level in those two species. These two species generally have highly similar protein orthologs with a median amino acid identity of 86% (calculated with http://ekhidna2.biocenter.helsinki.fi/AAI/ (Medlar *et al*. 2018), and both have the entire set of known recombination proteins. For example, RadA and RadB, of *H. volcanii* and *H. mediterranei* are 94% and 95% identical. However, Rad50 and Mre11 are surprisingly divergent, sharing only 82% and 78% identity, respectively. Thus, the difference between these species could lie at the level of resection of DNA by the Mre11-Rad50 complex, which should be tested experimentally by swapping these factors. Alternatively, a difference in RadB affinity for single-stranded DNA, or the differences in gene expression (as determined in our RNA-Seq data) between the two species could explain the differences described above.

Polyploidy confers an array of immediate advantages to the organism. However, in the absence of an accurate DNA segregation mechanism, the evolutionary long-term consequences of polyploidy, according to the logic of Muller’s ratchet (Muller 1964) and later models (Markov and Kaznacheev 2016) would lead to a high segregation load and the accumulation of deleterious recessive alleles, even resulting in giving rise to non-viable daughter cells. Therefore, a polyploid yet non-mitotic organism would face a strong purifying selection against the existence of the species. By extension, any extant polyploid organism has either found a way to escape the ratchet (Ludt and Soppa 2019) or at least substantially slow it down.

Several mechanisms have been suggested that can result in the organisms overcoming of the evolutionary trap of polyploidy. Interestingly at least half of these suggested strategies are coupled with or mechanistically connected to recombination [(Markov and Kaznacheev 2016)]: Ploidy cycle, namely the periodic decrease in polyploidy such that the daughter cells are subjected to selection, equalization of the chromosomes by gene-conversion thus exposing said deleterious alleles to selection in their homozygote state, lateral gene transfer and recombination that were shown to have a synergistic effect of rescuing otherwise ‘doomed’ populations by creating viable chromosomes from the fragments of existing genes, and acquired chromosomes the latter phenomenon potentially enhanced by cell-cell mating.

Here we show that while *H. volcanii* might escape the ratchet via efficient chromosomes equalization, *H. mediterranei* is much slower in doing so. According to the models, recombination is an important part of many ratchet evading mechanisms, but it shows a much higher rate in *H. volcanii* than in *H. mediterranei*. How *H. mediterranei* escapes Muller’s ratchet while still maintaining substantial “heterozygosity” for multiple generations remains to be elucidated.

## Materials and Methods

### *H. volcanii* and *H. mediterranei* strain construction

*H. volcanii* strains used in this study were derived from the parental strain H133 (Allers *et al*. 2004) that contains the deletions Δ*trpA* (HVO_0789, tryptophan auxotrophy) Δ*hdrB* (HVO_2919, thymidine auxotrophy) (table 1). H133 also contains deletions in the *leuB* and *pyrE2* genes that were not used in this work. Furthermore, deletion of the *crtI* (HVO_2528) gene, which is involved in biosynthesis of the carotenoid pigment bacterioruberin, and is not essential nor has an effect on fitness (Turkowyd *et al*. 2020) resulted in a white phenotype. Strain WR780 [Δ*trpA*, Δ*hdrB*, *1585::trpA* (red)] was constructed by replacing the non-essential ASC-like gene HVO_1585 in the Δ*crtI* strain with the *trpA* gene. Strain WR807 [Δ*trpA*, Δ*hdrB*, Δ*crtI 1585::hdrB* (white)] was constructed by replacing HVO_1585 with *hdrB*. Essentially the same process was repeated with *H. mediterranei*, but for practical reasons the *pyrE* gene (HFX_0318, uracil auxotrophy) was used as a selective marker instead of *folA* to generate strain UG482, along with *trpA* (HFX_0748) to generate strain UG480 This gene has been used before as a selectable marker in *H. mediterranei* (Naor *et al*. 2012)

### Culture conditions

Non-selective medium refers to Hv-YPC medium (Allers *et al*. 2004) supplemented with 50 μg/ml thymidine. Selective media were based on Hv-Ca medium (Allers *et al*. 2004) supplemented with 50 μg/ml of uracil, tryptophan, or thymidine, as needed. All growth steps were carried out at 45°C. Liquid cultures were grown in 2.5 ml of medium, with agitation at 180 rpm.

### Plasmid and primers

Plasmids used to create *H. volcanii* strains were constructed by joining ^~^500bp of each flanking sequence of the gene to be deleted into the suicide vector pTA131 (Allers *et al*. 2004) using specific restriction enzymes constructed on primers (table 4). When *trpA* or *hdrB* was used to replace the deleted gene, the replacement gene under the ferredoxin promoter (*P_fdx_*) was excised from pTA106 (*trpA*) or pTA187 (*hdrB*) (Altman-Price and Mevarech 2009) with restriction enzymes EcoRI and XbaI (NEB) and inserted by restriction and ligation between the flanking regions into pTA131 cut with the appropriate enzymes.

**Table 4.**
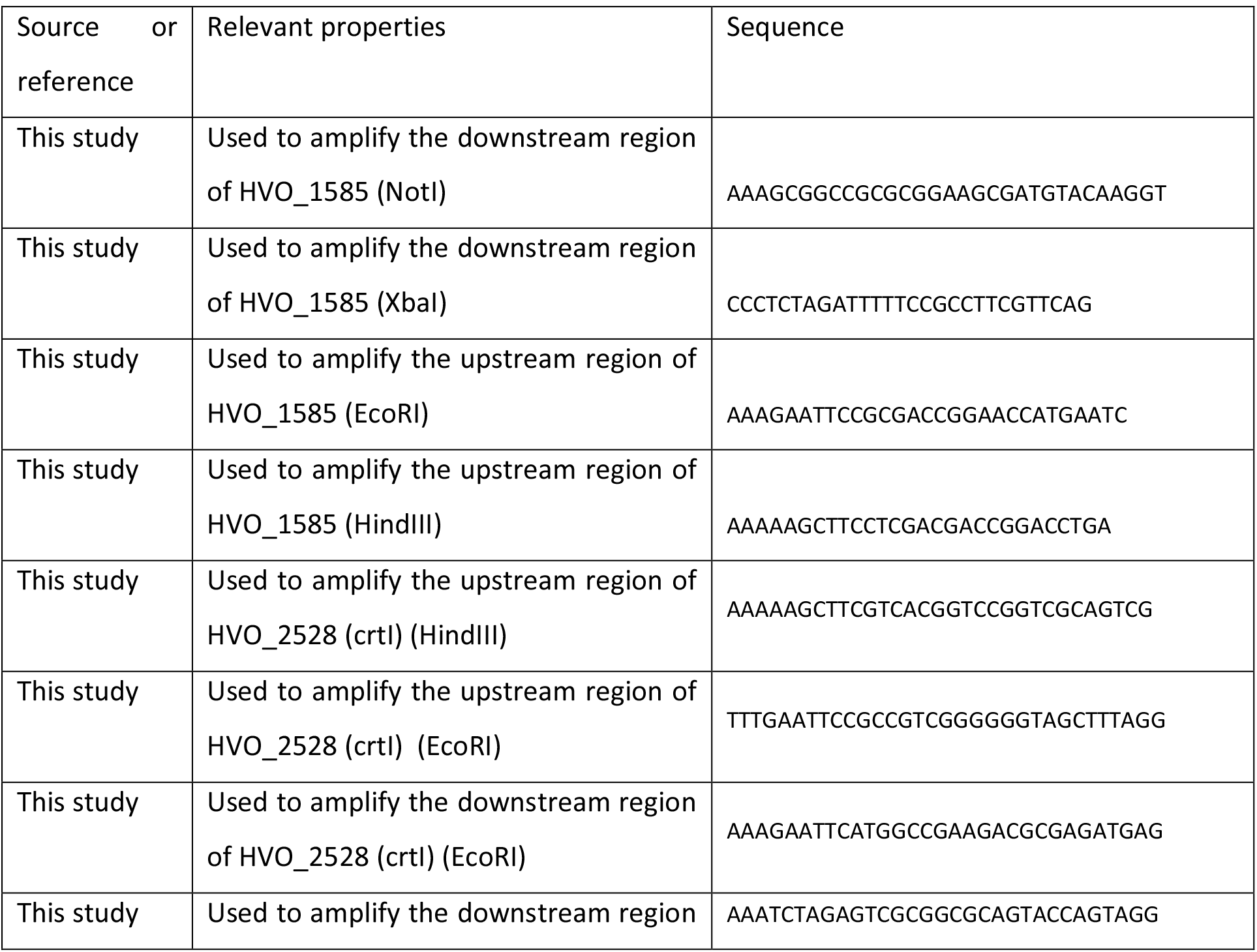
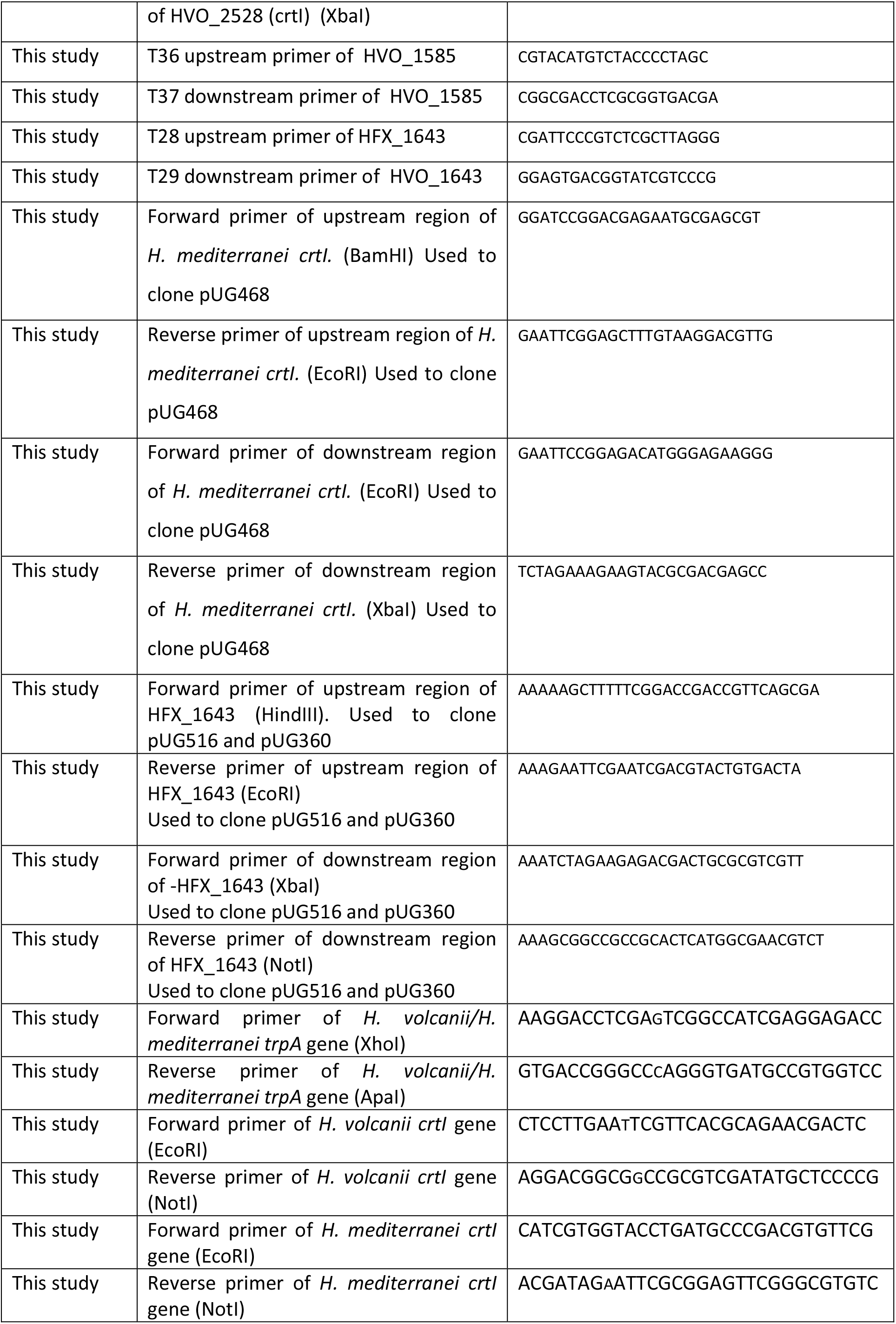
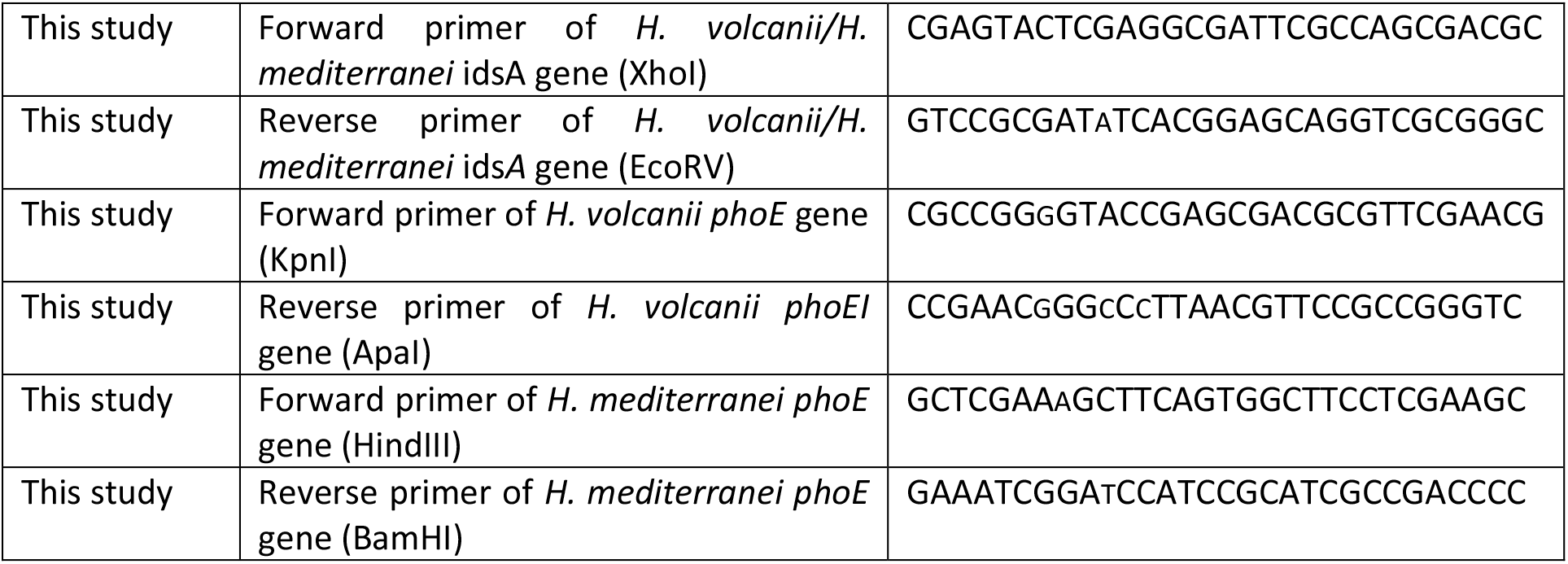
Primers used in this work

Plasmids used to create *H. mediterranei* strains were constructed using restriction enzymes or the Gibson-assembly method as described in table 5. Briefly, the sequences to be joined were amplified by PCR with ^~^25bp long overlapping regions on either side (table 4). ^~^30ng of each purified fragment were mixed and incubated with 10μl Gibson Assembly Master Mix (NEB) with pTA131 that was amplified with suitable primers, for 1 hour, in a final volume of 20μl. Ligated plasmids were transformed into *E. coli* (DH5α) using electroporation. Transformed bacteria were grown on LB-Agar plates supplemented with 100μg/ml ampicillin.

**Table 5.** Plasmids used in this work

| Source or reference | Relevant properties | Plasmid/strain |
| --- | --- | --- |
| (ALLERS *et al.* 2004) | Amp<sup>r</sup>; pBluescriptII with BamHI-XbaI fragment containing *P<sub>fdx</sub>-pyrE2* | pTA131 |
| Kindly provided by Thorsten Allers | Amp<sup>r</sup>; pUC19 containing *P<sub>fdx</sub>-trpA* | pTA106 |
| Kindly provided by Thorsten Allers | Amp<sup>r</sup>; pUC19 containing *P<sub>fdx</sub>-hdrB* | pTA187 |
| This study | pTA131 with *P<sub>fdx</sub>-hdrB* from pTA187 between HVO_1585 flanking sequences | pMM1172 |
| This study | pTA131 with *P<sub>fdx</sub>-trpA* from pTA106 between HVO_1585 flanking sequences | pMM1173 |
| This study | Flanking sequences of HVO_2528 (*crtI*) cloned into pTA131 using HindIII and XbaI | pMM1260 |
|  | Flanking sequences of *crtI* of *H. mediterranei* cloned into pTA131 using Gibson assembly | pUG468 |
| This study | pTA131 cut with HindIII and NotI and containing *P<sub>fdx</sub>-trpA* between *H. mediterranei* HFX_1643 flankings | pUG360 |
|  | pTA131 cut with HindIII and NotI and containing *P<sub>fdx</sub>-pyrE* between *H. mediterranei* HFX_1643 flankings | pUG516 |
| This study | Approximately 1000 bp of *H. volcanii* *trpA* gene cloned into pTA131 cut with XhoI and ApaI | pTA2686 |
| This study | Approximately 1000 bp of *H. volcanii* *crtI* gene cloned into pTA131 cut with XhoI and ApaI | pTA2690 |
| This study | Approximately 1000 bp of *H. volcanii* *idsA* gene cloned into pTA131 cut with XhoI and ApaI | pTA2694 |
| This study | Approximately 1000 bp of *H. volcanii* *phoE* gene cloned into pTA131 cut with XhoI and ApaI | pTA2698 |
| This study | Approximately 1000 bp of *H. mediterranei* *trpA* gene cloned into pTA131 cut with XhoI and ApaI | pTA2684 |
| This study | Approximately 1000 bp of *H. mediterranei* *crtI* gene cloned into pTA131 cut with XhoI and ApaI | pTA2688 |
| This study | Approximately 1000 bp of *H. mediterranei* *idsA* gene cloned into pTA131 cut with XhoI and ApaI | pTA2692 |
| This study | Approximately 1000 bp of *H. mediterranei* *phoE* gene cloned into pTA131 cut with XhoI and ApaI | pTA2696 |

All plasmids were sequenced to ensure they contained the desired inserts.

### Haloarchaeal strain construction

All transformations of archaea were performed using the PEG600 protocol described in the HaloHandbook (https://haloarchaea.com/halohandbook/). Deletion strains were constructed by transforming cells with pTA131 plasmid containing 500bp-long flanking regions of the targeted gene, and further selection steps as previously described (Bitan-Banin *et al*. 2003). Gene replacements were done by transforming archaea with approximately 2 μg of linear, purified PCR products composed of 500 bp-long flanking regions of the unwanted gene surrounding the gene to be inserted. Transformed colonies were then selected for by plating on Hv-Ca media supplemented with the appropriate metabolites. All strains were verified using PCR and selective plating.

### Mating protocol

Cultures were grown overnight and adjusted to an OD_600_ of 1. 2ml of each strain were mixed and filtered through a 0.2 μm filter using a syringe to dispose of excess media. The filter was placed on a non-selective plate and incubated overnight. Next, the cells were suspended in selective media and plated on selective plates. The resulting colonies were streaked on selective plates to ensure they contained both selective markers (see above).

### Heteropolyploids growth assay

Cultures were grown overnight and diluted to an OD_600_ of 1. A volume of 100 μL of each culture were used to inoculate 2 mL of rich medium. The culture was left to shake at 45 °C and samples were taken as described in the text. For the 1-week assays, 100 μL of the culture was used to inoculate a fresh 2 mL medium every 1-2 days.

### Pop-in assay

Plasmid-chromosome recombination assays, otherwise referred to here as “pop-in” assays, were performed as described in (Lestini *et al*. 2010). Transformations were plated on Hv-Ca and Hv-YPC plates to analyze the transformation efficiency and viable count, respectively. The number of colonies on both plates were counted and the recombination rate was calculated as a fraction of transformants on Hv-Ca plates compared to Hv-YPC.

### RNA isolation and RNA-seq analysis

*H. volcanii* and *H. mediterranei* cultures were grown to an OD650 ≈ 0.2. Cells were pelleted at 3,500 x *g* for 8 minutes and the pellet resuspended in 1 ml of Trizol (Invitrogen) and incubated for 5 minutes at room temperature. 200 μl chloroform was added and vortexed for 15 seconds and further incubated at room temperature for 3-5 minutes. Cell lysate was centrifuged at 4°C at 12,000 x *g* for 15 minutes. The top aqueous layer was transferred very carefully to a clean tube and ethanol precipitated. DNAase digestion was carried out using Ambion TURBO DNase as per manufacturer’s instructions. The integrity of the RNA was checked using PCR, agarose gel electrophoresis, Qubit and Agilent TapeStation. The RNA-Sequencing was carried out by the Deep Seq facility at the University of Nottingham. Sequencing and was performed using Illumina NextSeq500 instrument with a 150-cycle mid output sequencing kit to generate approx. 10million paired end reads per sample (2 × 75bp). *H. volcanii* DNA repair and replication homologues were obtained from (*Perez-Arnaiz et al*. 2020). *H. mediterranei* homologues were obtained using a BLASTN and SyntTax analysis. (Oberto 2013).

## Supplementary Figures

**Supplementary Figure 1.**
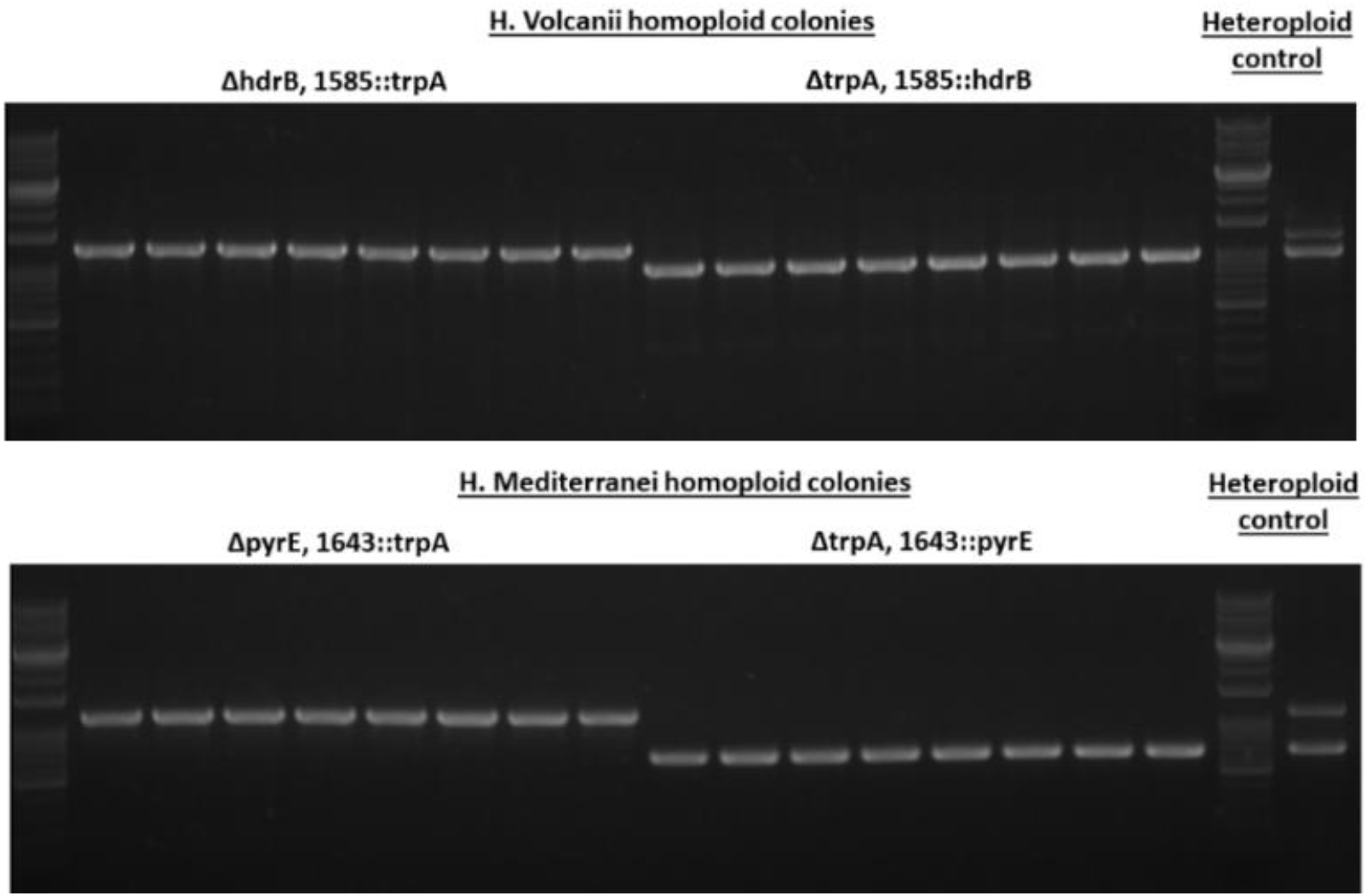
Similarity to parent strains among homopolyploid cells after mating and subsequent loss of heteropolyploidy. Representative PCR results. 5% of the colonies that were considered homopolyploid after streaking were also tested by PCR on locus 1585 of H. volcanii and 1643 of H. mediterranei. The results completely matched the observed phenotype.

**Supplementary Table 1.**
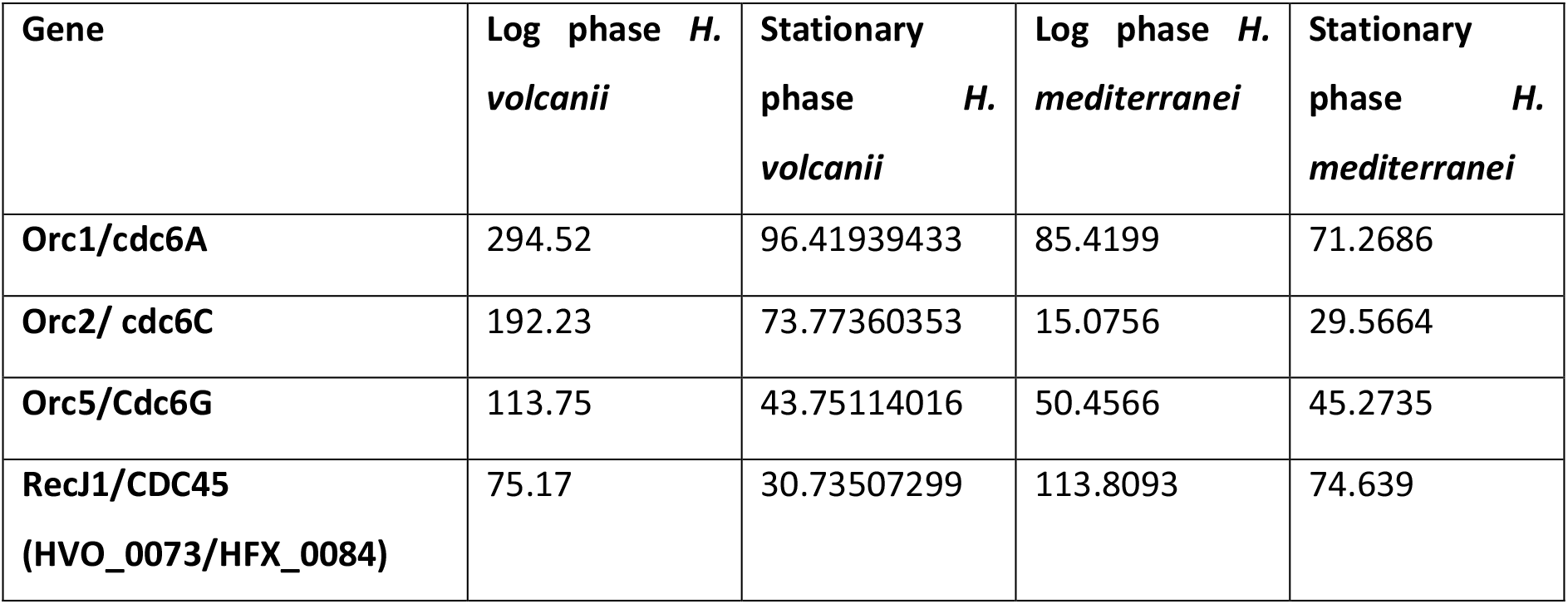
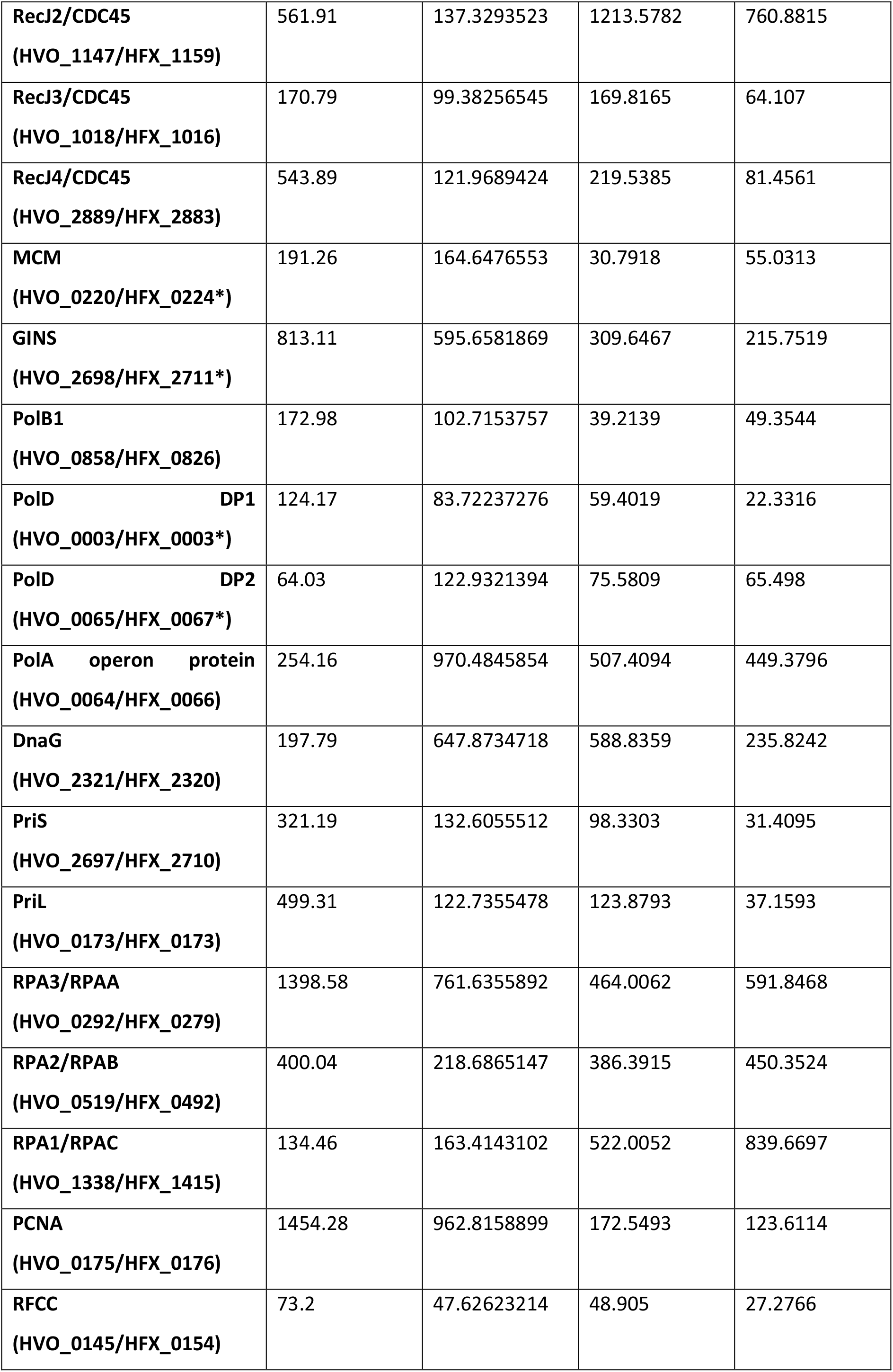
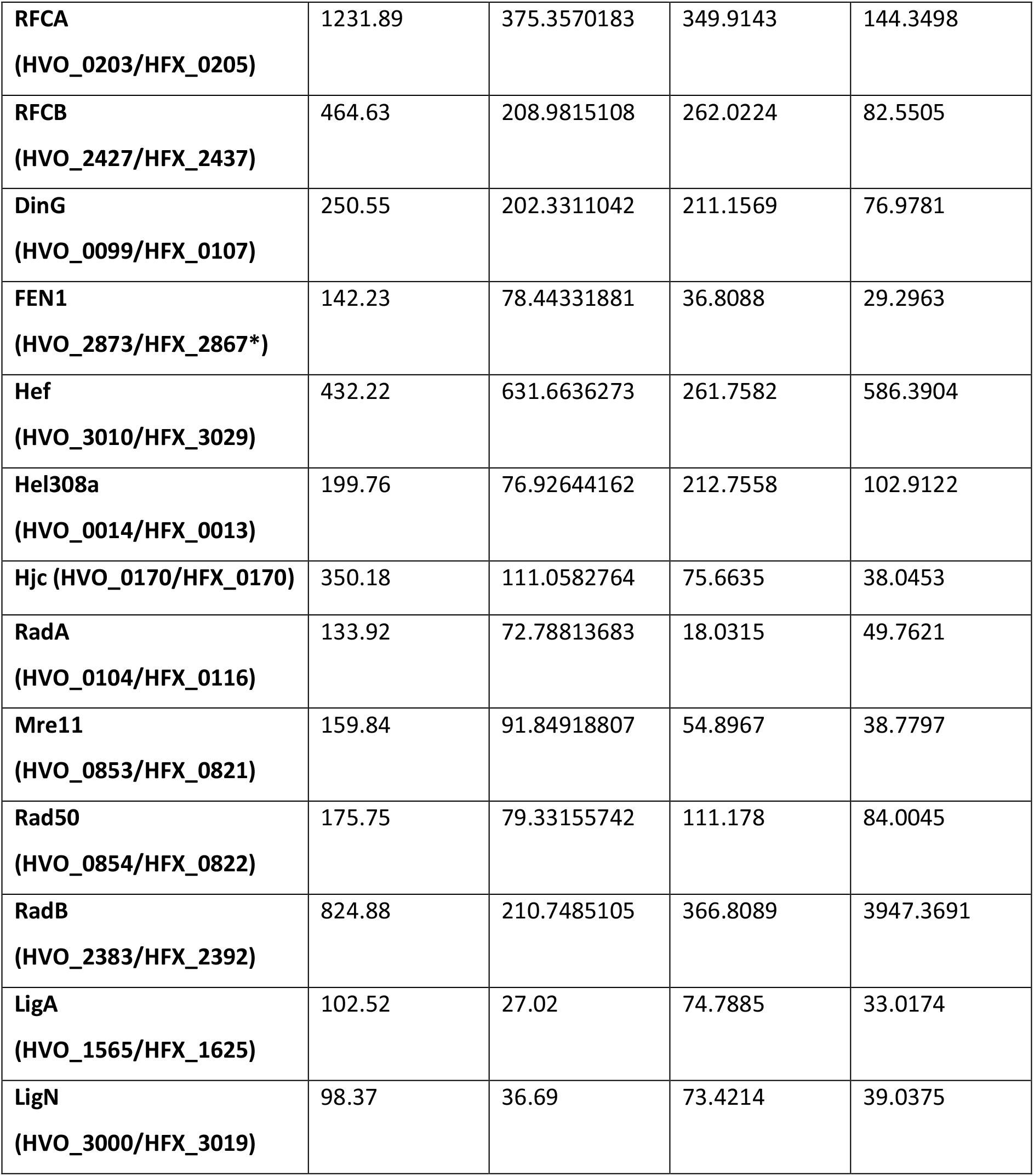
Comparing RNA expression between *H. volcanii* and *H. mediterranei*. Comparison of RNA expression levels of DNA replication and homologous recombination factors in *H. volcanii* and *H. mediterranei* as denoted by RPKM (Reads per Kb transcript per Million reads) values.

